# The dynamic interplay of PIP_2_ and ATP in the regulation of the K_ATP_ channel

**DOI:** 10.1101/2021.05.06.442933

**Authors:** Tanadet Pipatpolkai, Samuel G. Usher, Natascia Vedovato, Frances M Ashcroft, Phillip J. Stansfeld

## Abstract

ATP-sensitive potassium (K_ATP_) channels couple the intracellular ATP concentration to insulin secretion. K_ATP_ channel activity is inhibited by ATP binding to the Kir6.2 tetramer and activated by phosphatidylinositol-4,5-bisphosphate (PIP_2_). Here, we use molecular dynamics (MD) simulation, electrophysiology and fluorescence spectroscopy to show that ATP and PIP_2_ occupy different binding pockets that share a single amino acid residue, K39. When both ligands are present, K39 shows a greater preference to co-ordinate with PIP_2_ than ATP. A neonatal diabetes mutation at K39 (K39R) increases the number of hydrogen bonds formed between K39 and PIP_2_, reducing ATP inhibition. We also find direct effects on nucleotide binding from mutating E179, a residue proposed to interact with PIP_2_. Our work suggests PIP_2_ and ATP interact allosterically to regulate K_ATP_ channel activity.

## Introduction

Pancreatic ATP-sensitive potassium (K_ATP_) channels couple the energy status of the pancreatic beta-cell to insulin secretion^1^. Recent cryo-EM studies have provided high-resolution structures of the K_ATP_ channel complex, which comprises a central tetrameric pore formed from four inwardly rectifying potassium channel (Kir6.2) subunits, surrounded by four regulatory sulphonylurea receptor 1 (SUR1) subunits^2^. Binding of ATP to Kir6.2 closes the channel^3^, while binding of phosphoinositides, such as phosphatidylinositol-4,5-bisphosphate (PIP_2_), increases the channel open probability^4^. The ATP-binding site on Kir6.2 has been identified in several cryo-EM structures^5–10^. The PIP_2_ binding site has been resolved in structural studies of related Kir channels (Kir2.2, Kir3.2)^11,12^ but not yet for Kir6.2. However, its structure has been previously predicted using site-directed mutagenesis and coarse-grained molecular dynamics (CG-MD) simulation^13–15^.

Mutations in the K_ATP_ channel lead to diseases of insulin secretion. Channel hyperactivation is associated with reduced insulin secretion and neonatal diabetes mellitus (NDM) whereas reduced channel activity leads to enhanced insulin secretion and congenital hyperinsulinism (CHI)^16^. Previous studies have shown that many NDM mutations cluster at the ATP-binding site, disrupting the ATP-sensitivity of the channel^17^. Some NDM mutations also interfere with PIP_2_ regulation: for example E179K enhances PIP_2_ stimulation of the K_ATP_ current (as indicated by a reduction in neomycin block) and increases the predicted PIP_2_ binding affinity^15,18^.

The mechanism by which PIP_2_ reduces ATP inhibition of the channel has long been debated. Previous studies have shown that application of PIP_2_ markedly increases the channel open probability (P_open_) and reduces channel sensitivity to ATP^19^. Because an increased channel open probability is associated with reduced ATP inhibition^20,21^, it is possible that at least part of the effect of PIP_2_ is mediated via changes in P_open_. However, it has also been argued that PIP2 may have an additional effect on ATP sensitivity that is independent of Popen20. Both molecules carry similar negatively charged phosphate groups, and previous studies have proposed that PIP_2_ competes with ATP for the same binding site on the C-terminus of the protein^22^. However, comparison of recent structural studies of the channel with bound ATP^5,6^, and docking and molecular dynamics simulations with PIP_2_ suggest that ATP and PIP_2_ have separate binding pockets^23,24^.

In this study, we used atomistic molecular dynamic simulations (AT-MD) to determine the dynamics of K39 when both ATP and PIP_2_ occupy their respective binding sites. These data show that K39 can co-ordinate with both ATP and PIP_2_, but with stronger preference to PIP_2_ when both ligands are present. We used a combination of electrophysiology and fluorescence spectroscopy to assess how mutations which affect PIP_2_ binding modulate ATP inhibition and nucleotide binding. These support the simulation findings and suggest how the mutations give rise to clinical disease.

## Results

In this study, we used both atomistic molecular dynamic simulations and functional studies to explore the relationship between the ATP and PIP_2_ binding sites of Kir6.2.

### Simulation studies: PIP_2_ and ATP binding site on Kir6.2

We simulated the PIP_2_ and ATP-binding sites in the Kir6.2 tetramer in the absence of SUR1 in order to exclude ATP interactions with SUR1. Previous simulations have shown there is no difference in the PIP_2_ binding site when SUR1 is present^15^. Although the structure of the Kir6.2/SUR1 octameric complex has been resolved^5,6,10^, there is no structure of Kir6.2 in either a PIP_2_-bound or an ATP-bound conformation when SUR1 is absent. Thus, we built two atomistic simulation systems: Kir6.2 with ATP and Kir6.2 with PIP_2_, and simulated each for 380 ns. To ensure that the initial protein structure was stable after converting to an atomistic system, we used pairwise root mean square deviation (RMSD) analysis over the last 300ns of the simulation on the C_α_ atom of Kir6.2 as a measure of the stability of protein’s tertiary structure. We observed that the C_α_ RMSD never deviated more than 4 Å across the trajectory in all simulation set-ups. Interestingly, the simulations with bound ligand (PIP_2_ or ATP) showed slightly less C_α_ rearrangement than the apo state. This suggests that the 3D structure of the protein is highly stable throughout our simulation, and maybe further stabilised by the ligand (Figure 1, Supplementary figure 1). To investigate the effect of PIP_2_ and ATP binding at their binding sites, we select amino acid residues that were within 4 Å of the ligand for >40% of the time and defined them as contacting residues. We found that in the last 300 ns of the simulation, the contacts made with the ligands were in good agreement with previous electrophysiological and computational studies (Figure 1A,1B)^13,14^.

**Figure 1.**
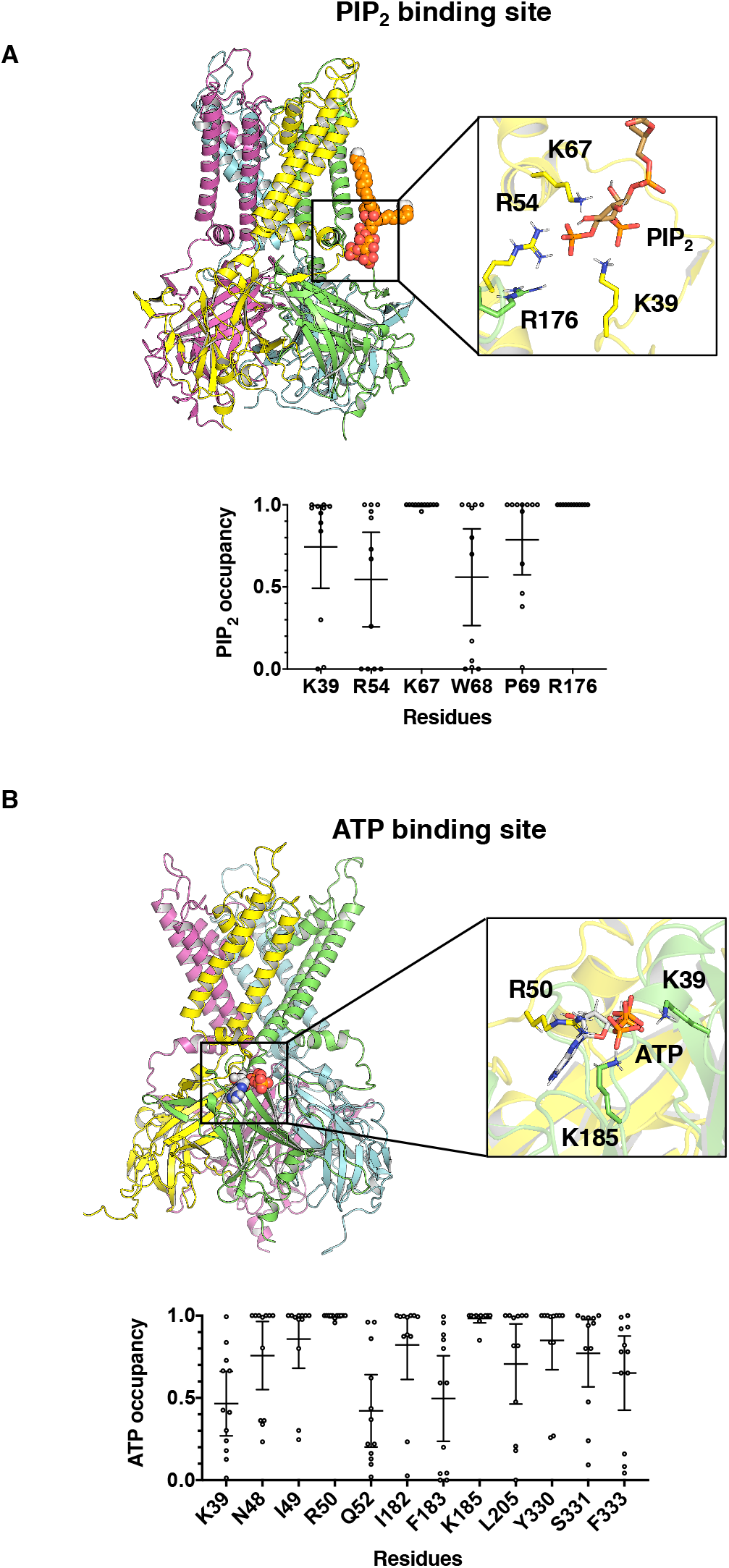
ATP and PIP_2_ binding sites. A) Above, structure of Kir6.2 tetramer with bound PIP_2_. Inset Interactions between PIP_2_ (bronze) and basic residues in two chains of Kir6.2 (yellow and green). Below, fraction of time that residues which contact the PIP_2_ headgroup >40% of the time in 4 subunits of three 300 ns simulations (a total of n = 12) lie in 4 Å proximity to the channel (PIP_2_ occupancy). The error bar indicates the 95% confidence interval around the mean. B) Above, Structure of Kir6.2 tetramer with bound ATP. Inset, interactions between ATP (CPK) and basic residues in two chains of Kir6.2 (yellow and green). Below, fraction of time that residues which contact ATP >40% of the time in all 4 subunits of three 300 ns simulations (a total of n = 12) lie in 4 Å proximity to the channel (ATP occupancy). The error bar indicates the 95% confidence interval around the mean.

In the PIP_2_ binding site, we found that R54 and K67 from one subunit, and R176 from an adjacent subunit co-ordinate with the 4’ phosphate of PIP_2_, and that K39 co-ordinates with the 5’ phosphate. Other uncharged residues (W68 and P69) that lie at the membrane-water interface also make strong contact with PIP_2_. With the exception of P69, mutations at these residues have previously been shown to alter channel ATP sensitivity, increase the open probability and/or alter PIP_2_ activation^24–27^. By evaluating the root mean square fluctuation (RMSF) of all contact residues, we found that the binding of PIP_2_ statistically reduces the dynamics of the K67 and R176 side chains (Supplementary figure 2A). However, only the difference at K67 showed biological significance (defined as a decrease of ~1 Å or more). These results agree well with previous coarse-grained simulations and hence, validate the co-ordination geometry of PIP_2_ in its binding site.^13,15^.

In the ATP binding site, we found that R50 co-ordinates with both the β and the γ phosphate, K39 co-ordinates with the γ phosphate and K185 co-ordinates with the α and the β phosphate (Figure 1B). Both R50 and K185 dynamics were stabilised when ATP binds to the channel (Supplementary figure 2B). These findings agree with previous studies in which the ATP-binding residues in Kir6.2 were mapped using site-directed mutagenesis ^28,29^. Interestingly, the sidechain of K39 which is *ca*. 7 Å from the ATP molecule in the cryo-EM structure moves towards ATP and makes a contact in some simulations. All residues that contact ATP in the cryo-EM structure of the octameric Kir6.2/SUR1 complex (N48, I49, Q52, I182, F183, L205, Y330, S331, F333) also make contact in our simulations. These residues have been found to be crucial for ATP inhibition in functional studies ^28^. Mutations in residues that make contact with ATP also cause neonatal diabetes ^17^.

### The competition between PIP_2_ and ATP for K39 co-ordination

Previous studies have shown that PIP_2_ reduces channel ATP inhibition^4,19,30,31^. However, it was not clear if PIP_2_ competes directly for the ATP binding site or if it interferes with ATP-dependent gating (or both). Comparison of the cryo-EM structure of the ATP binding site with the predicted PIP_2_ binding site suggest they lie *ca*. 25 Å from one another^5,15^. In our studies, K39 contacts both ATP and PIP_2_ in independent simulations, but the position of the amine head group is different. We next explored the dynamics of K39 when both ligands occupied their respective binding sites (Figure 2A). We calculated the distance between ATP or PIP_2_ and the amine group of K39. We observed that the position of the amine head group of K39 oscillated between co-ordination with ATP and PIP_2_, but favoured PIP_2_ more strongly (Figure 2B and Supplementary figure 3). Interestingly, a significant decrease (P = 0.001) in ATP contact only occurred at residue K39 when PIP_2_ was present (Figure 2C). Thus, we hypothesise that K39 may change its co-ordination from ATP to PIP_2_ when both molecules are bound. Interestingly, the contact between K39 and PIP_2_ remained unchanged in both in the presence and the absence of ATP (Figure 2D). Thus, we propose that the salt bridges between K39 and ATP are broken in the presence of the PIP_2_, causing the amine group on K39 to swing towards the PIP_2_ headgroup.

**Figure 2.**
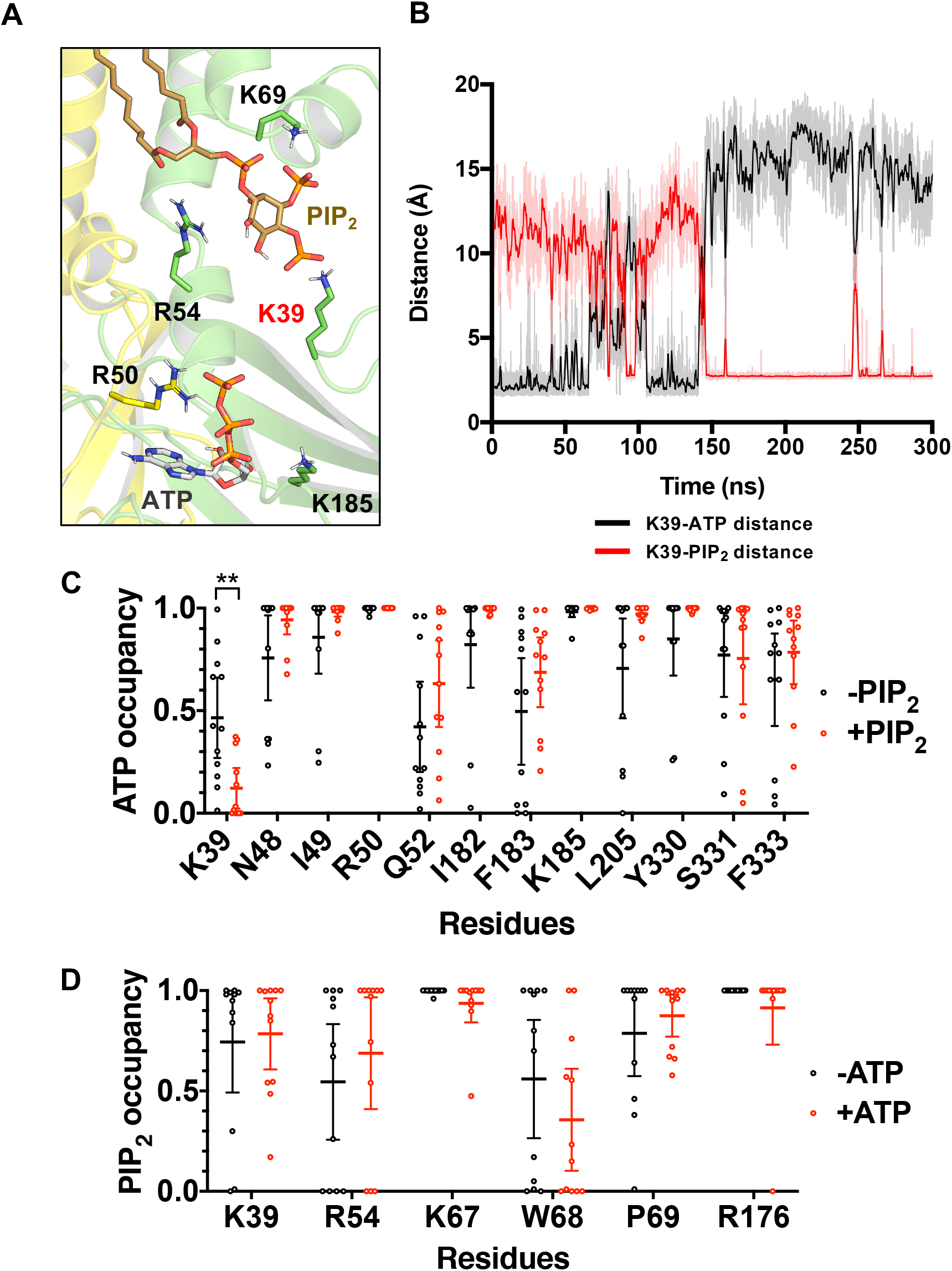
Changes in the ATP and PIP_2_ binding sites when both ligands are present. A) Interactions between Kir6.2 (two subunits are indicated in yellow and green), ATP (CPK) and the PIP_2_ (bronze) headgroup. Only the basic residues of the protein are shown. B) A representative trace showing the distance calculation between K39 and ATP (black) and K39 and PIP_2_ (red) across a single 300ns trajectory. The darker lines show running averages of every 1 ns of the simulation. C) ATP contact analysis showing the fraction of time that residues which contact the ATP molecule >40% in all 12 repeats of 300 ns simulations are in 4 Å proximity to the channel (ATP occupancy) in the absence (black) and presence (red) of PIP_2_. The error bar indicates the 95% confidence interval around the mean. ** p< 0.01 using Student’s t-test. D) PIP_2_ contact analysis showing the fraction of time that residues which contact the PIP_2_ molecule >40% in all 12 repeats of 300 ns simulations are in 4 Å proximity to the channel (PIP_2_ occupancy) in the absence (black) and presence (red) of ATP. The error bar indicates the 95% confidence interval around the mean.

**Figure 3.**
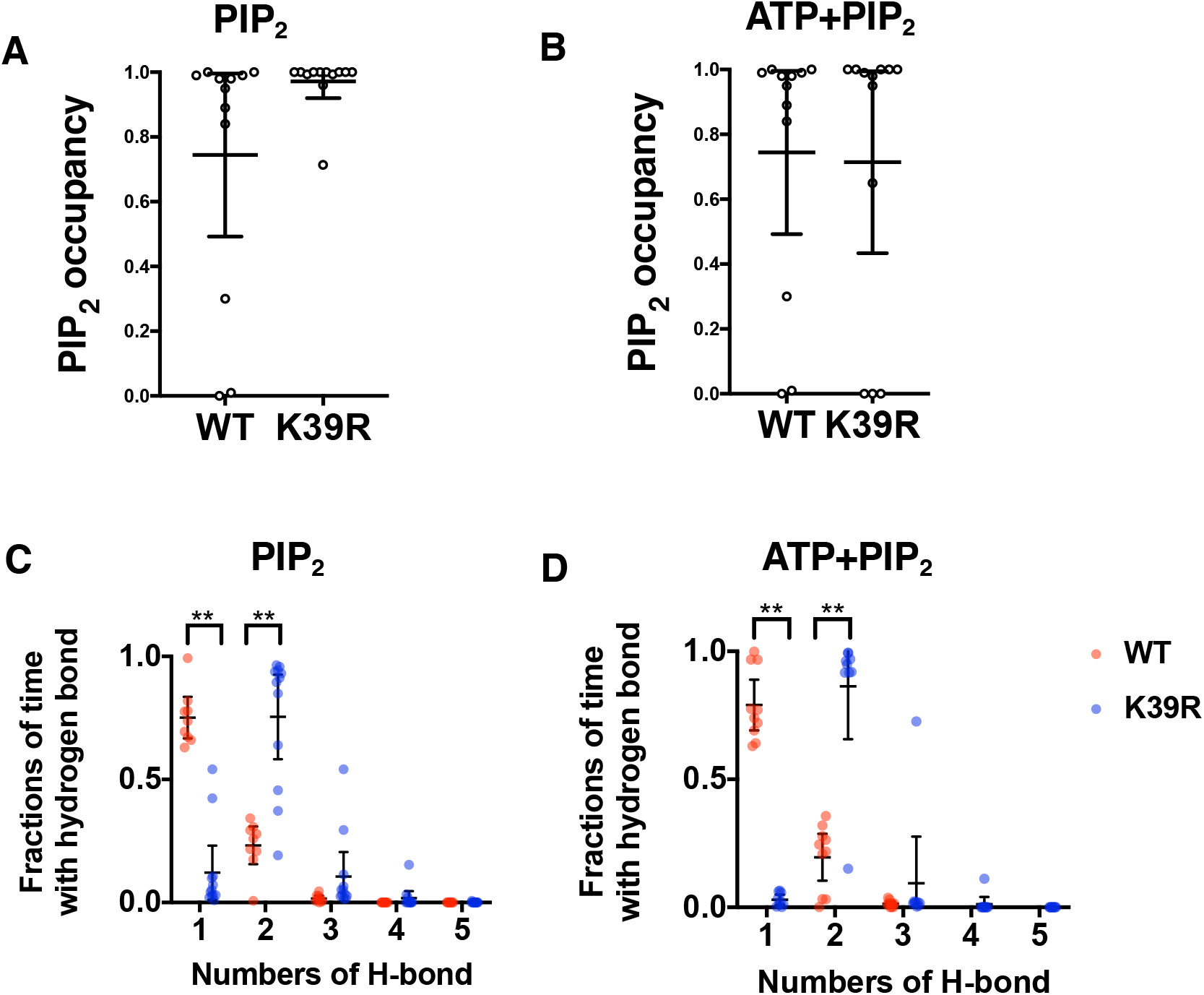
The K39R mutation changes the PIP_2_ binding configuration. A, B) PIP_2_ contact analysis showing the fraction of time that K39 (or K39R) is within 4 Å proximity to PIP_2_ across all four subunits in three 300 ns simulations (a total of n = 12). The error bar indicates the 95% confidence interval around the mean in (A) the absence of PIP_2_ or (B) the presence of PIP_2_. C) Hydrogen bond analysis showing the fraction of time when K39 (red) or K39R (blue) forms a different number of H-bonds with the PIP_2_ headgroup in 4 subunits of three 300 ns simulations (a total of n = 12), when only PIP_2_ is present. The error bar indicates the 95% confidence interval around the mean. ** p < 0.01 (Student’s t-test). D) Hydrogen bond analysis showing the fraction of time when K39 (red) or K39R (blue) forms different numbers of H-bond with the PIP_2_ headgroup in 4 subunits of three 300 ns simulations (a total of n = 12) when both ATP and PIP_2_ are present. The error bar indicates the 95% confidence interval around the mean. ** p < 0.01 (Student’s t-test).

With the exception of K39, no other residues in either the ATP or PIP_2_ binding site changed their contact probabilities, when both ligands were simultaneously present in their respective binding sites.

### Neonatal diabetes mutation (K39R)

A mutation at K39 (K39R) is associated with transient neonatal diabetes^32^. This mutation does not alter the charge (as both lysine and arginine are positively charged), but an amine is replaced with a guanidium group. This would likely increase the channel affinity to both PIP_2_ and ATP. We simulated Kir6.2 containing the K39R substitution and compared the contacts between the guanidium group of the arginine and PIP_2_ or ATP. In simulations with ATP alone, residue 39 in the mutant channel (arginine) spent more time in contact with ATP than that (lysine) of the wild-type channel (Supplementary figure 4). In simulations with PIP_2_ alone, residue 39 in both wild-type and mutant channels spent almost all of its time co-ordinated with PIP_2_, but the 95% confidence interval was small for the mutant channel. When both ATP and PIP_2_ were present, we found that the contact probability of residue 39 with the PIP_2_ headgroup was not significantly different in the either absence or the presence of the ATP (Figure 3A, 3B). From this, we conclude that K39R mutation does not affect channel preference for PIP_2_ in either the presence or the absence of ATP.

To quantify the strength of the interaction between PIP_2_ and K39 or K39R, we carried out a hydrogen bond (H-bond) analysis to determine the number of H-bonds formed between the PIP_2_ headgroup and the side chain of residue 39. Note that arginine can form up to 5 H-bonds spread over 3 amine groups, whereas lysine is only able to form a maximum of 3 bonds from a single amine. We observed that the guanidium group on the arginine forms two hydrogen bonds with the PIP_2_ headgroup, whereas the lysine amine group forms only a single hydrogen bond. In both cases, the hydrogen bonds were formed with the 5’ phosphate on the PIP_2_ inositol headgroup both in the presence and the absence of the ATP (Figure 3C, 3D). This suggests that the K39R mutation enhances the strength of the interaction of residue 39 with PIP_2_. We postulate this leads to reduced channel inhibition by ATP, and thereby impairs insulin secretion leading to neonatal diabetes.

### Electrophysiology and Fluorescence Spectroscopy

We next used functional studies to determine if the predicted change in PIP_2_ binding associated with the K39R mutation affects the ATP sensitivity of the K_ATP_ channel and if this is mediated by changes in ATP binding.

There are two distinct hypotheses for how PIP_2_ binding to Kir6.2 affects nucleotide binding to Kir6.2. These are non-exclusive and both may occur simultaneously. The first (theory A) postulates that PIP_2_ binding preferentially stabilises the open state of the channel, increasing the open probability (P_open_) and antagonising ATP inhibition indirectly. This theory predicts that ATP binding *per se* will be unaffected but this has not been measured directly. However, it has previously been shown that PIP_2_ increases P_open_ ^4,30^ and that an increase in P_open_ reduces ATP inhibition of the K_ATP_ channel^20,21^. The second (theory B) is that PIP_2_ binding to the channel antagonises ATP binding directly: this could be described as a local allosteric effect. In the case of K39, our simulations are consistent with theory B, as K39 appears to coordinate both ATP and PIP_2_, and therefore binding of one necessarily will have a local allosteric effect on binding of the other.

To test this hypothesis, we examined the functional effect of mutations at K39 in two ways; by measuring the ability of ATP to inhibit the K_ATP_ current and by exploring ATP binding using a FRET-based assay with fluorescent trinitrophenyl (TNP)-ATP as a congener for ATP^33,34^. The latter method utilises FRET between channels labelled with the fluorescent unnatural amino acid 3-(6-acetylnaphthalen-2-ylamino)-2-aminopropanoic acid (ANAP) at residue W311 of Kir6.2 and TNP-ATP. Incorporation of ANAP into Kir6.2 was achieved as described previously^33–36^. A GFP tag was also added at the C-terminus of Kir6.2 to facilitate identification of transfected cells. HEK293T cells were transfected with control or mutant Kir6.2 and wild-type SUR1 and currents measured in inside-out membrane patches. All constructs described in this section have GFP at the C-terminus and are co-expressed with SUR1 (unless otherwise stated) to increase surface membrane trafficking^37^. Experiments were carried out in the absence of Mg^2+^, to prevent Mg-nucleotide modulation of channel activity by SUR1. Constructs labelled with ANAP at residue W311 are denoted as Kir6.2*.

A reduction in K_ATP_ channel inhibition by ATP may be mediated by an increase in the unliganded P_open_, by a decrease in ATP binding, and/or by modifying the transduction process by which ATP occupancy of its binding site is translated into altered gating. Because K39 forms part of both PIP_2_ and ATP binding pockets, we also examined a residue (E179) that is predicted to form part of the PIP_2_ binding site but which does not contact ATP in the cryo-EM structure^5–7,9,10,13,15^. We might therefore expect that any effect on ATP binding and inhibition caused by E179 mutations would be mediated indirectly, either via changes in P_open_ or by allosteric effects on binding.

### Reduction in K_ATP_ current inhibition by ATP and TNP-ATP

We first examined the effect of mutations at K39 and E179 on ATP inhibition. We introduced three different residues at K39 (K39A, K39E, K39R) and two different residues at E179 (E179A, E179K). When these mutations were introduced into Kir6.2, we observed a statistically significant reduction in current inhibition by ATP (i.e. an increase in the IC_50_) for each mutation tested, with the exception of K39A, which was indistinguishable from the control construct (Supplementary figure 5 and Supplementary table 1). Labelling the nucleotide binding site of Kir6.2 with ANAP (Kir6.2*) caused a reduction in ATP inhibition (from an IC_50_ of 34 µM to 112 µM). However, all mutations that caused a significant decrease in ATP inhibition in the Kir6.2 background still caused a significant decrease when in the Kir6.2* background (Figure 4A-D, and Supplementary table 2).

**Figure 4.**
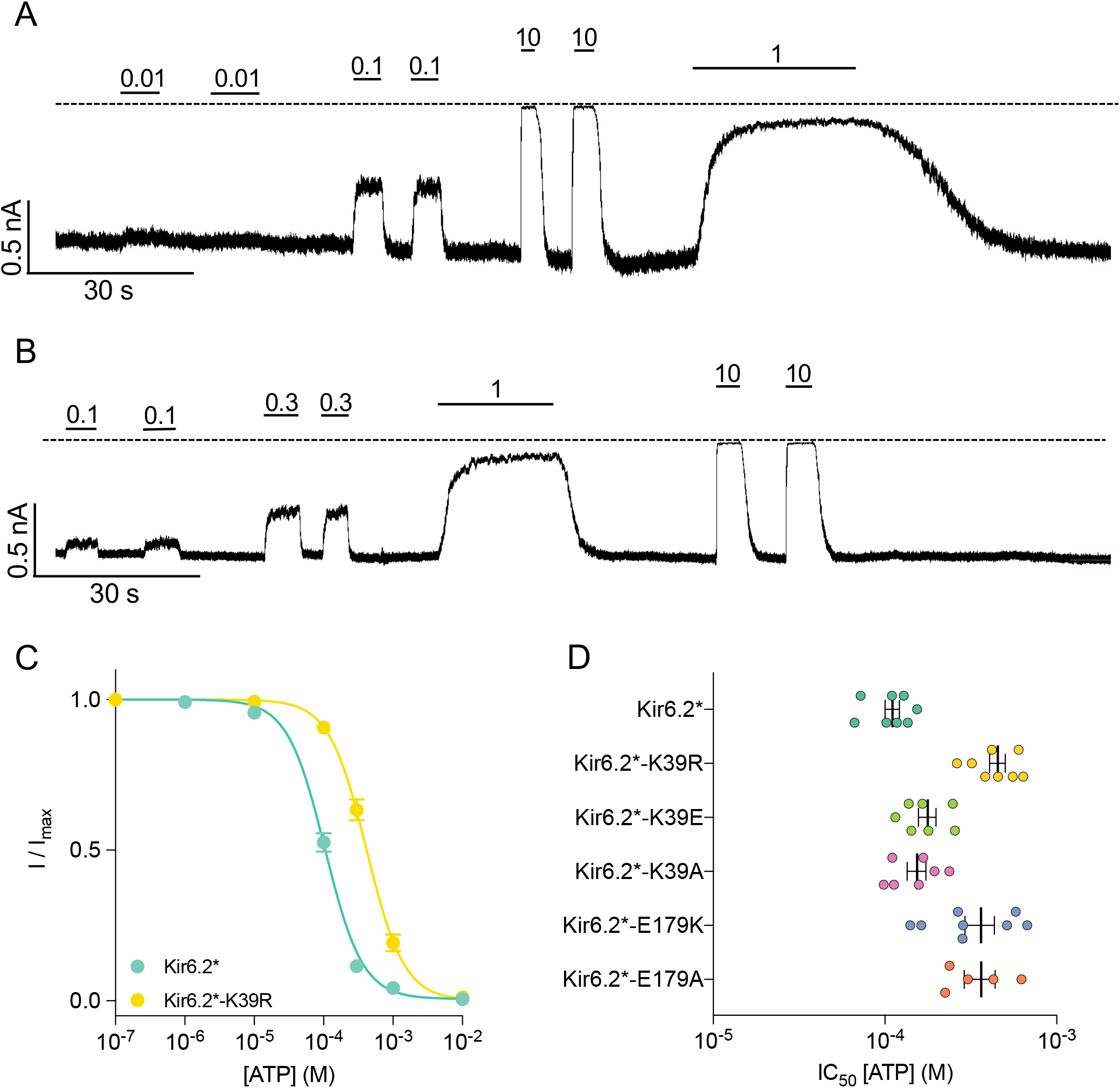
Effect of ATP on K_ATP_ current. A, B) Representative traces for Kir6.2* (A), and Kir6.2*-K39R (B). Top solid bars mark the different ATP concentrations applied (in mM). Dashed line indicates the zero-current level. B) Concentration-response relationships for ATP inhibition of Kir6.2* and Kir6.2*-K39R currents (I/I_max_), fitted to the Hill equation. IC_50_: 105 µM for Kir6.2*, 417 µM for Kir6.2*-K39R, fitted with common, but free, *h*=1.6 amd I_max_ fixed to 1. C) IC_50_s from ATP current inhibition derived from Hill fits to individual experiments are plotted as coloured data points. The means and 95% confidence intervals for each construct are overlaid as black error bars. Each construct is labelled with GFP at the C-terminus, with ANAP at residue W311, and is co-expressed with SUR1.

As the simulations were conducted with Kir6.2 in the absence of SUR1, we also examined ATP inhibition when mutant Kir6.2 was expressed alone (Supplementary table 3). We observed low current amplitudes for all constructs except Kir6.2-K39A, for which we were unable to resolve any current. The IC_50_ values for each construct were dramatically increased (by roughly an order of magnitude) when compared to the values measured when co-expressed with SUR1. Importantly, the mutant channels remained less sensitive to ATP than controls. We also transfected mutant Kir6.2* alone, but were unable to resolve any macroscopic currents. Thus, we focused our experiments on Kir6.2/SUR1 constructs.

Because measurement of ATP binding requires the use of fluorescent TNP-ATP, we next measured the effects of the five mutations introduced into Kir6.2 on TNP-ATP inhibition of the channel (Figure 5A-D). TNP-ATP is a more potent inhibitor of the K_ATP_ channel than ATP ^33,34^, reducing the IC_50_ from 34 µM to 1.4 µM (Supplementary table 4). Four of the five mutations (K39A, K39E, E179A, E179K) displayed a similar reduction in current inhibition when tested with TNP-ATP to that found for ATP (Figure 5G, Supplementary table 4).

**Figure 5.**
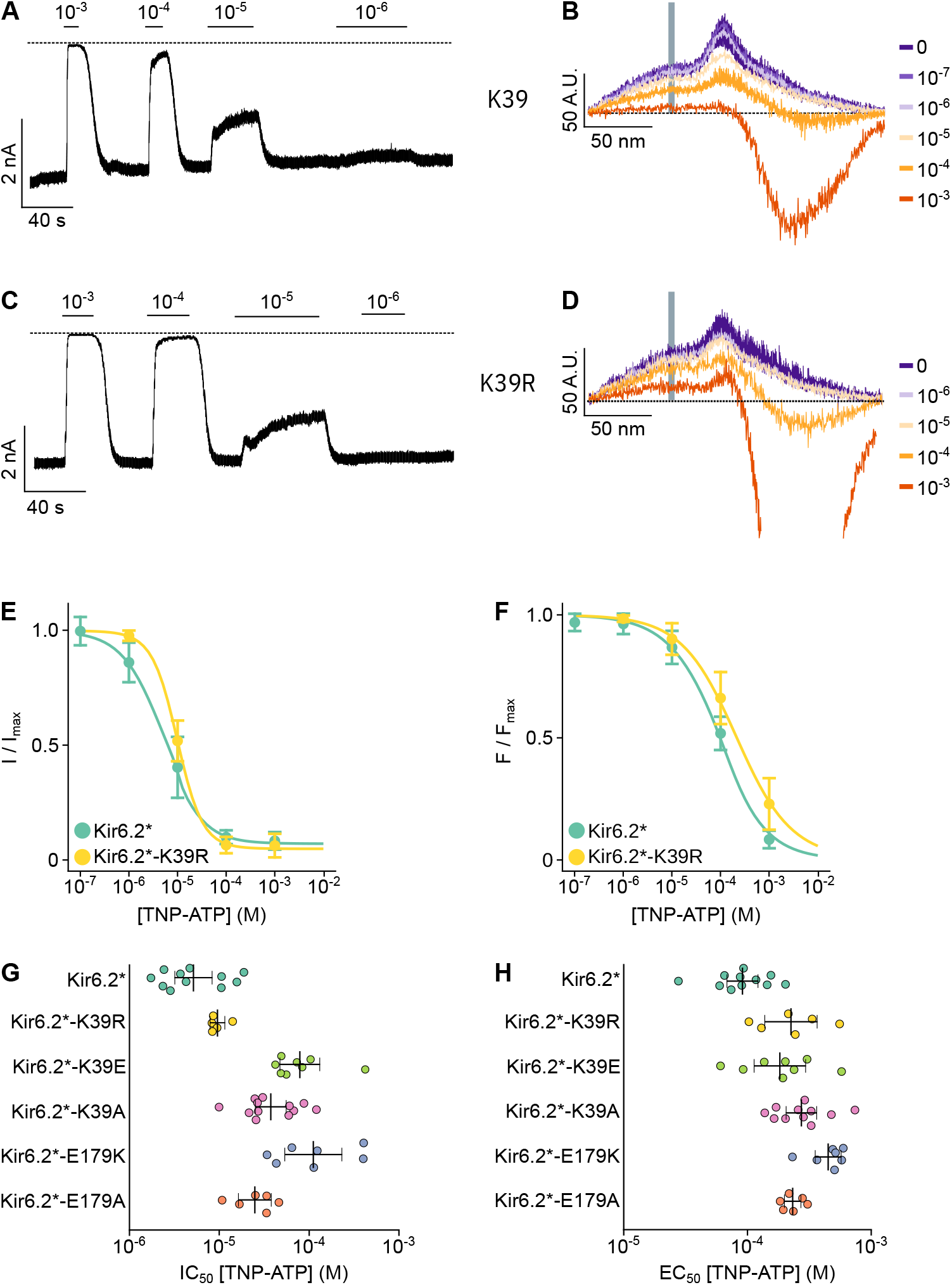
Effect of TNP-ATP on K_ATP_ current. A, B) Representative current (A) and fluorescence (B) traces for Kir6.2*. A) Top solid bars indicate application of TNP-ATP, with concentrations given in M. Dashed line indicates the zero-current level. B) The grey bar indicates the fluorescence peak corresponding to ANAP. The colours indicate application of TNP-ATP, with concentrations given in M. The dashed line indicates the zero-fluorescence level with respect to background. C, D) Representative current (C) and fluorescence (D) traces for Kir6.2*-K39R. E, F) Concentration-response relationships for TNP-ATP current inhibition (C, I/I_max_) and fluorescence quenching (D, F/ F_max_) of Kir6.2* (Kir6.2*, green) and Kir6.2*-K39R (Kir6.2*-K39R, yellow). Data points are the mean and error bars are the 95% confidence intervals. Smooth lines are fits to the Hill equation, with coefficients given in figure 5, table supplement 1. G, H) IC_50_s from TNP-ATP current inhibition (G) and EC_50_s from TNP-ATP fluorescence quenching (H) derived from Hill fits to individual experiments are plotted as coloured data points. The means and 95% confidence intervals for each construct are overlaid as black error bars. Each construct is labelled with GFP at the C-terminus, ANAP at residue W311 and co-expressed with SUR1.

Finally, we simultaneously measured current inhibition and nucleotide binding by TNP-ATP in the Kir6.2* background using patch-clamp fluorometry (Figure 5E,F,H and Supplementary table 5,6). Both E179A and E179K mutations resulted in a significant reduction in current inhibition (from an IC_50_ of 7µM for Kir6.2* to 28 µM and 169 µM for Kir6.2*-E179A and Kir6.2*-E179K respectively) and TNP-ATP binding (from an EC_50_ of 94 µM to 170 µM and 171 µM for Kir6.2*-E179A and Kir6.2*-E179K respectively). However, despite observing a significant reduction in TNP-ATP binding to Kir6.2*-K39R (from an EC_50_ of 94 µM to 209 µM), we did not find a significant decrease in current inhibition.

We fit our combined TNP-ATP binding and inhibition data to an allosteric model of K_ATP_ described previously^33,34^. This model seeks to describe channel gating with three parameters: *L* is a measure of the unliganded channel open probability, which in part reflects PIP_2_ regulation, *K*_*A*_ is the binding affinity for nucleotides, and *D* is the global allosteric constant which represents the degree to which nucleotide binding favours the closed state of the channel. We estimated posterior probability distributions for these parameters for each mutation (Supplementary figure 6A-B) using a Bayesian Markov chain Monte Carlo (MCMC) method^38^. We observed very broad posterior probabilities for *L*, which we believe reflects a range of observed open probabilities, likely due to differences in channel rundown or variability in membrane PIP_2_ between patches. However, this does not affect our estimates of the other two parameters *D* and *K*_*A*_ ^33^.

Our parameter estimates for Kir6.2*-E179A and Kir6.2*-E179K channels suggest that the observed reduction in current inhibition by nucleotides is mostly due to a reduction in their ability to bind, as represented by decreases in the posterior probability distributions for *K*_*A*_ (Supplementary figure 6C). The E179K mutation also leads to a modest increase in *D*, implying that transduction between the ATP binding site and gating is weakened by the mutation. Each of the mutations at K39 leads to a reduction in *K*_A_, with the magnitude of the shift increasing R<E<A. In addition, K39E leads to a similar increase in *D* to that observed for E179K.

## Discussion

Our work provides a mechanistic explanation for how PIP_2_ and ATP binding influence one another to modulate the K_ATP_ channel activity. We show that in addition to its effects on P_open_ ^4,19,30^, PIP_2_ modulates ATP binding via long distance (E179) and local (K39) interactions.

The PIP_2_ binding site is conserved across Kir channel structures^11,12^ and previous simulation studies have suggested that residue E179 forms part of the PIP_2_ binding site of Kir6.2^13,15^. Mutation of E179 reduces neomycin sensitivity of KATP currents supporting a role for this residue in PIP_2_ binding^15^. Although E179 does not lie within the ATP-binding site^5,6,9,10^, the E179A and E179K mutations cause a significant decrease in current inhibition by ATP or TNP-ATP, both for the control Kir6.2 construct and for that labelled with ANAP. Binding of TNP-ATP was also significantly decreased by the E179A and E179K mutations. This is consistent with the idea that mutations at E179 influence ATP binding via changes in PIP_2_ binding or changes in P_open_. Analysis of these data using our MWC model indicates this cannot be due to a change in unliganded P_open_ alone, or solely due to a change in transduction, but rather due to a clear decrease in the microscopic *K*_*A*_. While this might be caused by a direct influence of the mutation at the ATP-binding pocket, this would not be consistent with the existing cryo-EM structures^5–7,9,10^. Thus, our data suggest that in the case of the E179 mutations, TNP-ATP binding is affected allosterically. Coupled with previous simulations showing that the E179K mutation enhances PIP_2_ binding^15^. We suggest that this allosteric effect may be mediated by a change in PIP_2_ binding.

Our atomistic MD simulations suggest that a single key residue, K39, forms part of both the ATP and PIP_2_ binding sites. When both ligands are present, K39 has a stronger preference for co-ordination with PIP_2_ than with ATP. This reduces the channel affinity for the ATP as K39 is no longer available to co-ordinate the ATP γ phosphate (Figure 6A-C). No other residue (in either site) alters its contact probability in the presence of the other ligand. The simulation data are supported by our finding that mutation of lysine 39 to arginine (K39R) led to a significant decrease in current inhibition by ATP. A similar reduction was found for K39E but not for K39A. However, a reduction in ATP sensitivity has been previously reported for the K39A mutation^26,39^. Introducing an ANAP label into Kir6.2 reduced ATP inhibition for control (K39) channels and even more for mutant (K39E/R/A) channels. Unfortunately, the effect of K39 mutations on TNP-ATP inhibition did not replicate the effects seen for ATP or in our simulations. In particular, there was no longer a significant difference in TNP-ATP inhibition when K39 was mutated to arginine. This may be due to the location of K39 in the ATP binding site – there is potentially a steric clash between the side chain of residue 39 and the TNP moiety of TNP-ATP (Supplementary figure 7). Although we saw shifts in the EC_50_ for fluorescence quenching, and associated shifts in the estimated microscopic association constant (*K*_*A*_) for TNP-ATP, it is difficult to interpret these shifts meaningfully. This is because differences in TNP-ATP binding may not accurately reflect differences in ATP binding for the K39 mutant channels.

**Figure 6.**
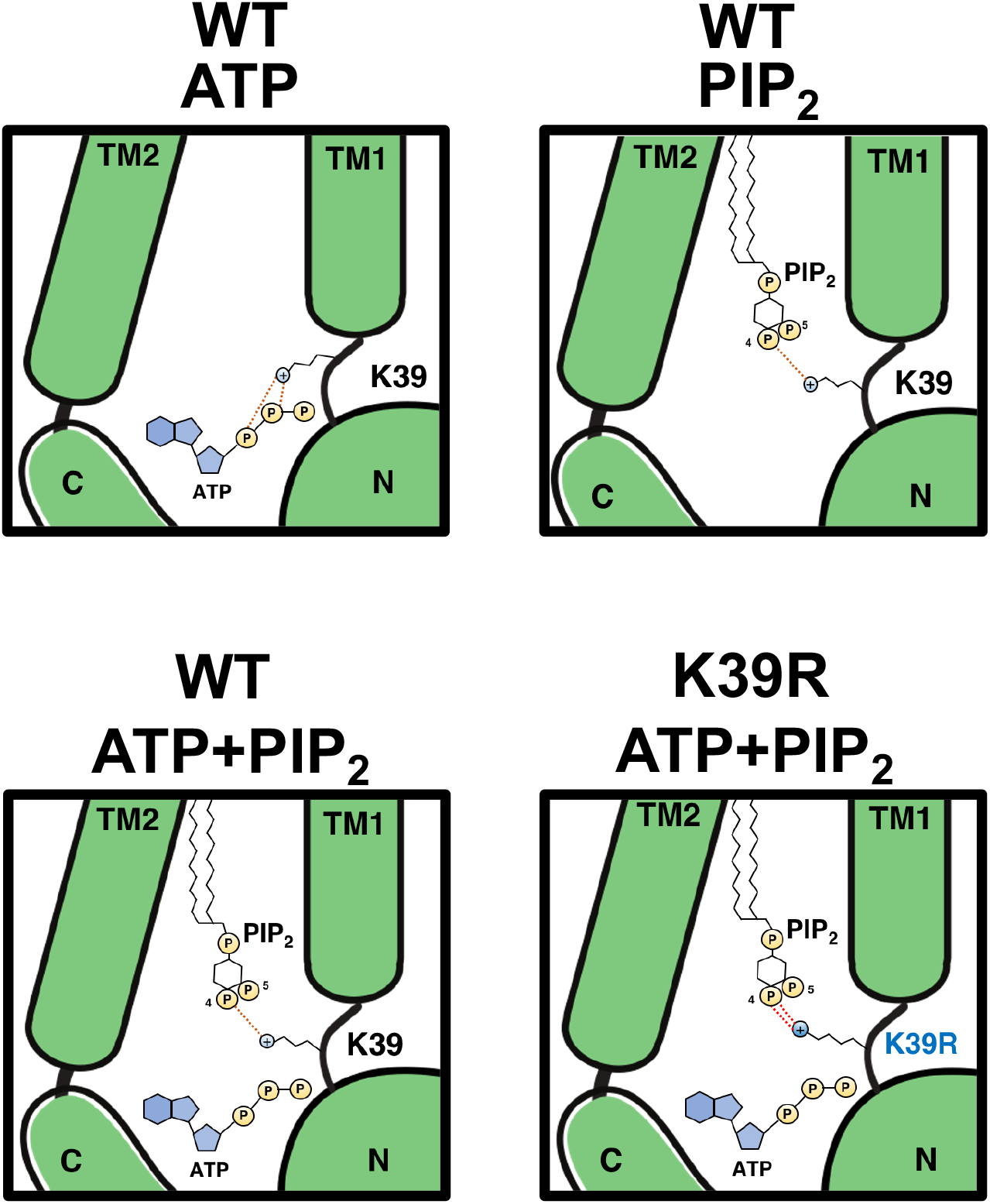
Schematic representation of the interaction between Kir6.2, ATP and PIP_2_. Interaction between ATP (silver sphere), PIP_2_ (hexagon with silver phosphate group – P) and Kir6.2 (green). A positive charge on K39 is denoted in yellow (or red in the K39R mutant). The orange dashed line represents hydrogen bonds between K39 and the ligand.

Nevertheless, the effects of the K39R mutation on current inhibition by ATP, together with our simulations, lead us to propose a mechanism for how the K39R mutation may cause transient neonatal diabetes. We found that the mutation increases the number of H-bonds between the side chain of residue 39 and the PIP_2_ headgroup (Figure 6D). This enhances PIP_2_ affinity, which in turn reduces the ability of ATP to close the channel and causes a reduction in insulin secretion that leads to neonatal diabetes.

In summary, our work is consistent with PIP_2_ regulating the function of Kir6.2 both by enhancing the P_open_ of the channel, and by allosterically modulating the binding of ATP. In addition, we have demonstrated a mechanism by which mutations at residue 39 can lead to neonatal diabetes. They do so by enhancing PIP_2_ binding which leads to a reduced sensitivity of Kir6.2 to ATP inhibition.

## Materials and methods

### Coarse-Grained system preparation

The human Kir6.2 model without SUR1 (PDB entry: 6BAA)^5^ was converted to a coarse-grained model using *martinize*.*py*, embedded in the palmitoyl oleoyl phosphatidylcholine (POPC) bilayer and solvated in water and 0.15M NaCl using a self-assembly MEMPROTMD pipeline^40^. All simulations were carried out in MARTINI2.2 biomolecular forcefield^41^. The tertiary and quaternary structures of the protein were maintained through the application of an elastic network with a force constant of 1,000 kJ/mol/nm^2^ between two coarse-grained backbone particles within 0.5-0.9 nm. A temperature of 323 K was maintained with V-rescale temperature coupling^42^, while 1 atm pressure was controlled using semi-isotropic Parrinello-Rahman pressure coupling^43^. The position of the coarse-grained PIP_2_ is taken from the chicken Kir2.2-PIP_2_:diC8^11^ after conversion to a coarse grain model^44^. Systems were energy minimised using the steepest descents algorithm and equilibrated for 1 μs. All simulations were carried out using GROMACS-2019.4^45^.

### Atomistic simulation set up

The coarse-grained simulation system (Kir6.2, lipids - PIP_2_ and POPC, ions and water) was converted to atomistic using the CG2AT pipeline^46^. The K39R mutant model was generated using PyMOL with the minimum initial stearic clashes^47^. The exact position of the ATP molecule was taken from the cryo-EM structure (PDB entry: 6BAA) and placed in the Kir6.2 ATP binding site. The initial position of the hydrogen atoms was added using PyMOL. All simulations were carried out using CHARMM36 biomolecular forcefield with the virtual sites on the proteins and lipids CH_3_ and NH_3_^+^ groups^48,49^, allowing the integration timestep of 4 fs in the production run. The force field for the ATP molecule was derived from the CHARMM-GUI^50,51^. In this study, 4 different conditions were set (Apo, ATP-bound, PIP_2_-bound, both ATP and PIP_2_ bound) and simulations were carried out for 3 repeats.

The systems were energy minimised using the steepest descents algorithm with non-hydrogen atoms restrained at 1000 kJ/mol/nm^2^. This was then followed by a 5 ns equilibration for the system where the C_α_ backbone on the protein and the non-hydrogen atoms on the ATP molecules were restrained with 1000 kJ/mol/nm^2^ with 4 fs timesteps. A temperature of 310 K was maintained with V-rescale temperature coupling, while 1 atm pressure was controlled using semi-isotropic Parrinello-Rahman pressure coupling^43^. The simulation was then further equilibrated with only C_α_ restraint on the protein for another 15 ns in similar conditions. Then a 380ns production run was implemented, in which the first 80 ns of the simulations were discarded as equilibration. All simulations, H-bond calculations and distance calculation were calculated using GROMACS-2019.4^45^. We used MDAnalysis to calculate pairwise RMSD to define change in protein dynamics across the trajectory^52–55^. As Kir6.2 is a symmetrical homo-tetramer, we assume that all sites behave identically under a short simulation timescale (380 ns). Thus, the results from our calculations are obtained over all 4 monomeric sites, making a total of 12 data points for all calculations.

### Molecular biology

Human Kir6.2 and SUR1 were subcloned into pCGFP_EU (GFP-tagged constructs) and pcDNA4/TO, respectively. Site-directed mutagenesis and amber stop codons were introduced using the QuikChange XL system (Stratagene; San Diego, CA), and verified by sequencing (DNA Sequencing and Services; Dundee, Scotland), as previously described^33^. HEK-293T cells were grown in in Dulbecco’s Modified Eagle Medium (DMEM, Sigma) with the addition of 10% fetal bovine serum, 100 U/mL penicillin, and 100 µg/mL streptomycin (Thermo Fisher Scientific; Waltham, MA) at 37°C, 5%/95% CO_2_/air. Cells were seeded in T25 flasks 24 hours, before transfection with TransIT-LT1 (Mirus Bio LLC; Madison, WI). Transfected cells were incubated at 33 °C, 5%/95% CO_2_/air. Protein expression and trafficking to the plasma membrane were optimized by including 300 μM tolbutamide in the transfection media (Yan et al., 2007; Lin et al., 2015). ANAP-tagged Kir6.2 constructs were cultured in the presence of 20 μM ANAP (free acid, AsisChem; Waltham, MA) and 48 hours post-transfection, cells were re-plated onto either poly-D-lysine coated glass-bottomed FluoroDishes (FD35-PDL-100, World Precision Instruments) or poly-L-lysine coated 35 mm petri dishes (Corning). pCDNA4/TO and pANAP were obtained from Addgene. peRF1-E55D (*Homo sapiens*) and pCGFP_EU (*Aequorea victoria*) were kind gifts from the Chin Laboratory (MRC Laboratory of Molecular Biology, Cambridge UK) and Gouaux Laboratory (Vollum Institute, Portland OR USA), respectively.

### Electrophysiology

Extracellular (pipette) solutions contained (in mM) 140 KCl, 1 EGTA, 10 HEPES (pH 7.3 with KOH). The intracellular (bath) solutions contained (in mM) 140 KCl, 1 EDTA, 1 EGTA, 10 HEPES (pH 7.3 with KOH). Inside-out patches were excised from transfected HEK cells using borosilicate glass pipettes (GC150F-15, Harvard Apparatus; Holliston, MA) pulled to a resistance of 1-3 MΩ. Data were acquired at a holding potential of −60 mV using an Axopatch 200B amplifier and a Digidata 1322A digitizer run through pClamp 9 software (Molecular Devices; San Jose, CA). Currents were low-pass filtered at 1-5 kHz and digitized at 10-20 kHz.

Patches were perfused with an 8-channel µFlow or a manual gravity perfusion system. Different concentrations of adenosine triphosphate (ATP, Sigma) or fluorescent trinitrophenyl adenosine triphosphate (TNP-ATP, Jena Bioscience) were added to the bath solution to assess current inhibition and/or nucleotide binding. Nucleotide-induced inhibition was corrected for rundown by alternating test concentrations of nucleotide solution with nucleotide-free solution. The inhibition was expressed as a fraction of the control currents before and after the test solution as described previously^56^. For experiments with TNP-ATP, the zero current level was determined by perfusing 10 mM BaCl_2_ at the end of each recording at a holding potential of +60 mV. Current inhibition data were fitted with the following Hill equation:

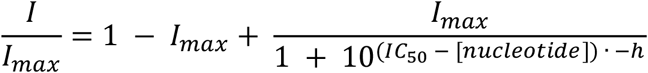

### Fluorescence measurements

Fluorescence spectra from excised patches were collected and analysed as described previously^33^. Briefly, the tip of the patch pipette was centred on the slit of the spectrometer immediately after patch excision. ANAP was excited using a 385 nm LED source (ThorLabs; Newton, NJ) with a 390/18 nm band-pass excitation filter, after which the emitted light passed through a 400 nm long-pass emission filter (ThorLabs) and an IsoPlane 160 Spectrometer (Princeton Instruments; Trenton, NJ) with a 300 grooves mm^-1^ grating. Images were collected with 1 s exposures on a Pixis 400BR_eXcelon CCD (Princeton Instruments). Spectra were corrected for background fluorescence, then ANAP intensity was calculated by averaging the fluorescence intensity measured between 469.5 nm and 474.5 nm. This intensity was corrected for bleaching with a single exponential decay. Fluorescence quenching data were fitted with the following Hill equation:

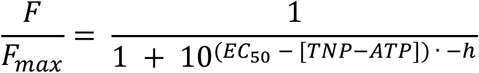

### Statistics and data presentation

Concentration-response data are plotted as the mean response at each tested concentration, with error bars representing the 95% confidence interval around the mean calculated from a normal distribution. IC_50_ and EC_50_ values obtained from patch-clamp and patch-clamp fluorometry experiments with TNP-ATP and ATP were compared with a one-way ANOVA, and differences between mutations at K39 and E179 to the control constructs were tested for significance with Dunnett’s post-hoc test.

### Monod-Wyman-Changeaux model fitting

The MWC-type model fitted here is described by the following sets of equations:

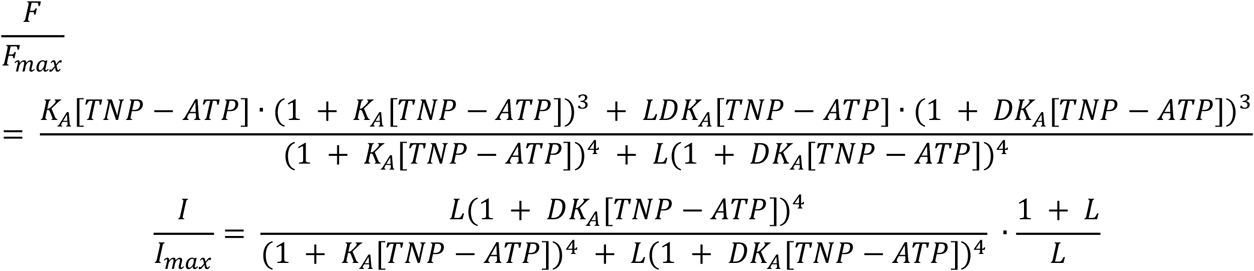

The full rationale behind this model choice is laid out in Puljung *et al* (2019). In this model, *L* represents an equilibrium constant where the K_ATP_ open probability (P_open_) is equal to *L*/(*L+1*), reflecting the ability of KATP to open and close in the absence of nucleotides. Each ligand binding event (*K*_*A*_) is independent and each bound ligand favours the closed state by the same factor (*D*). This model is fit to the combined current inhibition and fluorescence quenching data using the brms (Bayesian Regression Models using ‘ Stan’) package^57^ in R. Prior probability distributions were supplied for each parameter as follows:

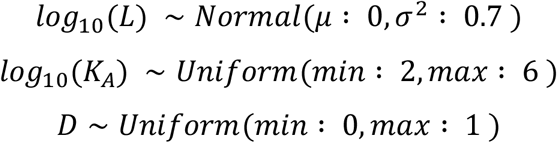

The model was run with tetrameric Kir6.2 and ATP (and /or PIP_2_) present in all 4 subunits for 4,000 iterations including a burn-in period of 2,000 iterations for a total of 8,000 samples. Each model parameter achieved a minimum effective sample size of 5,000 and a potential scale reduction statistic 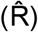 of 1.00.

## Acknowledgements

We thank Michael C. Puljung for discussions about the MWC modelling and interpretations of the parameters. We also thank Irfan Alibay, Robin Corey, Michael Horrell, Raul Terron-Exposito, and Owen Vickery for advice and technical support.

## Author contributions

T.P. performed the coarse-grained and atomistic molecular dynamics simulations. N.V. and S.U. performed the molecular biology, electrophysiology and fluorescence measurements. All authors jointly designed the experiments, analysed the data and wrote the manuscript.

---

**Supplementary figure 1.**
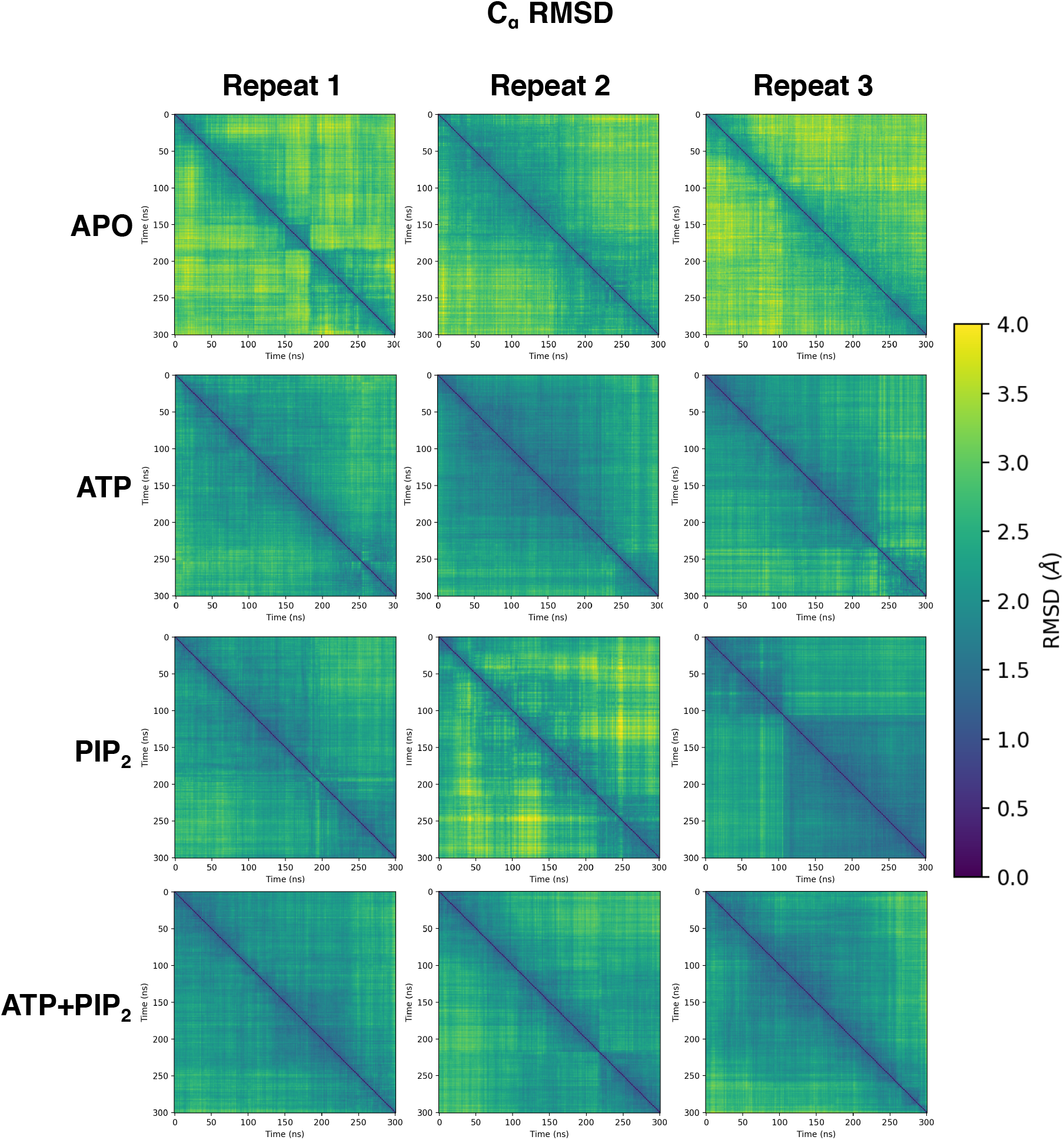
Overall protein dynamics in different states of the simulation. Pairwise Root Mean Square Deviation (RMSD) analysis of the C_α_ atoms of Kir6.2 in the different ligand-bound states, measured over the final 300 ns simulation trajectory (n=3). The colorbar indicates the RMSD value ranging from 0 to 4 Å.

**Supplementary figure 2.**
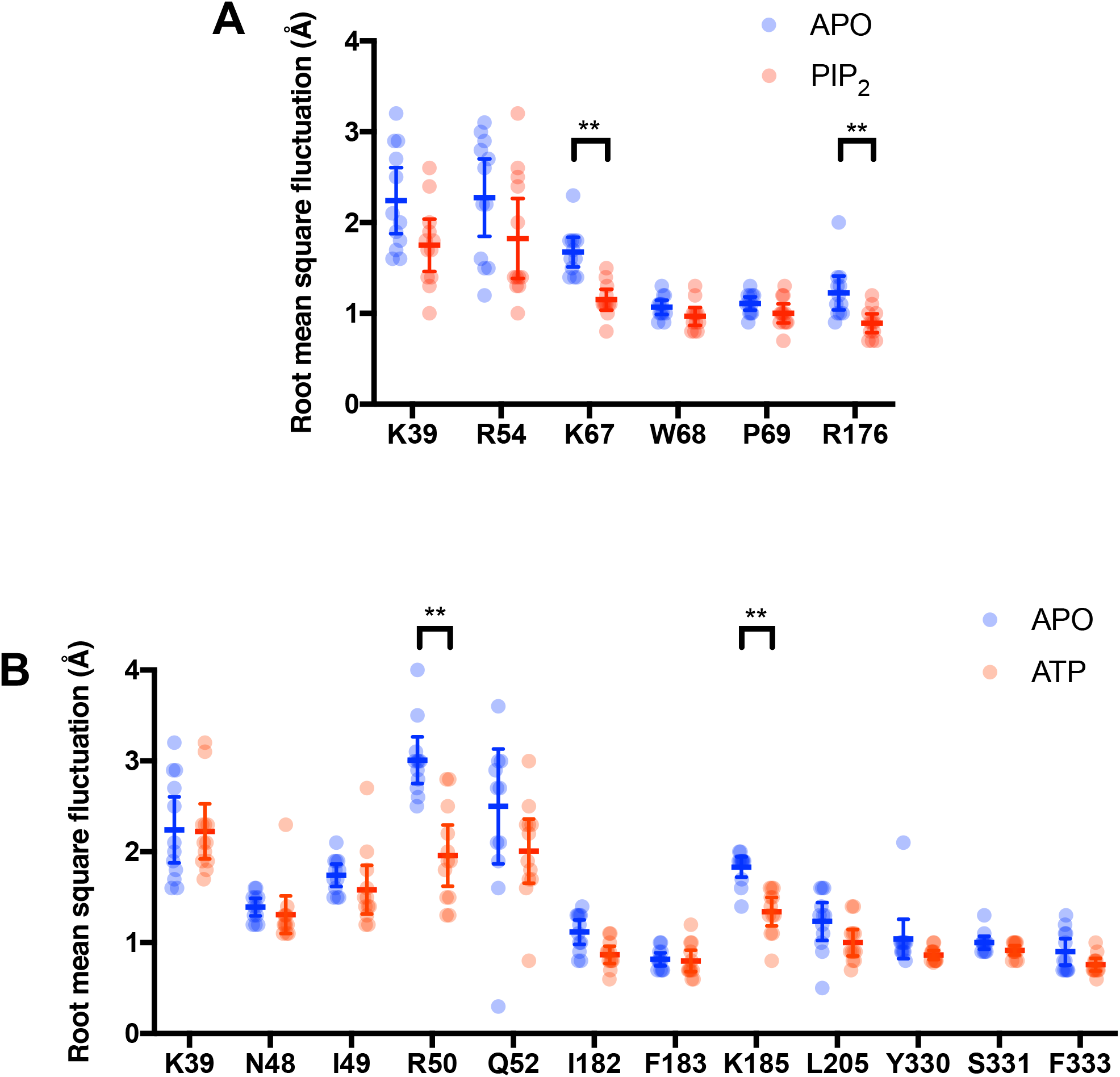
The dynamics of individual side chains in the simulation. A) Root mean square fluctuation (RMSF) analysis of the residues on the Kir6.2 tetramer which made contact with the PIP_2_ molecule in the absence (blue) and the presence (red) of PIP_2_ (n = 12). The error bar indicates the 95% confidence interval around the mean. ** p < 0.01 (Student’s t-test). B) Root mean square fluctuation (RMSF) analysis of the residues on the Kir6.2 tetramer which made contact with the ATP molecule in the absence (blue) and the presence (red) of ATP (n = 12). The error bar indicates the 95% confidence interval around the mean. ** p < 0.01 (Student’s t-test).

**Supplementary figure 3.**
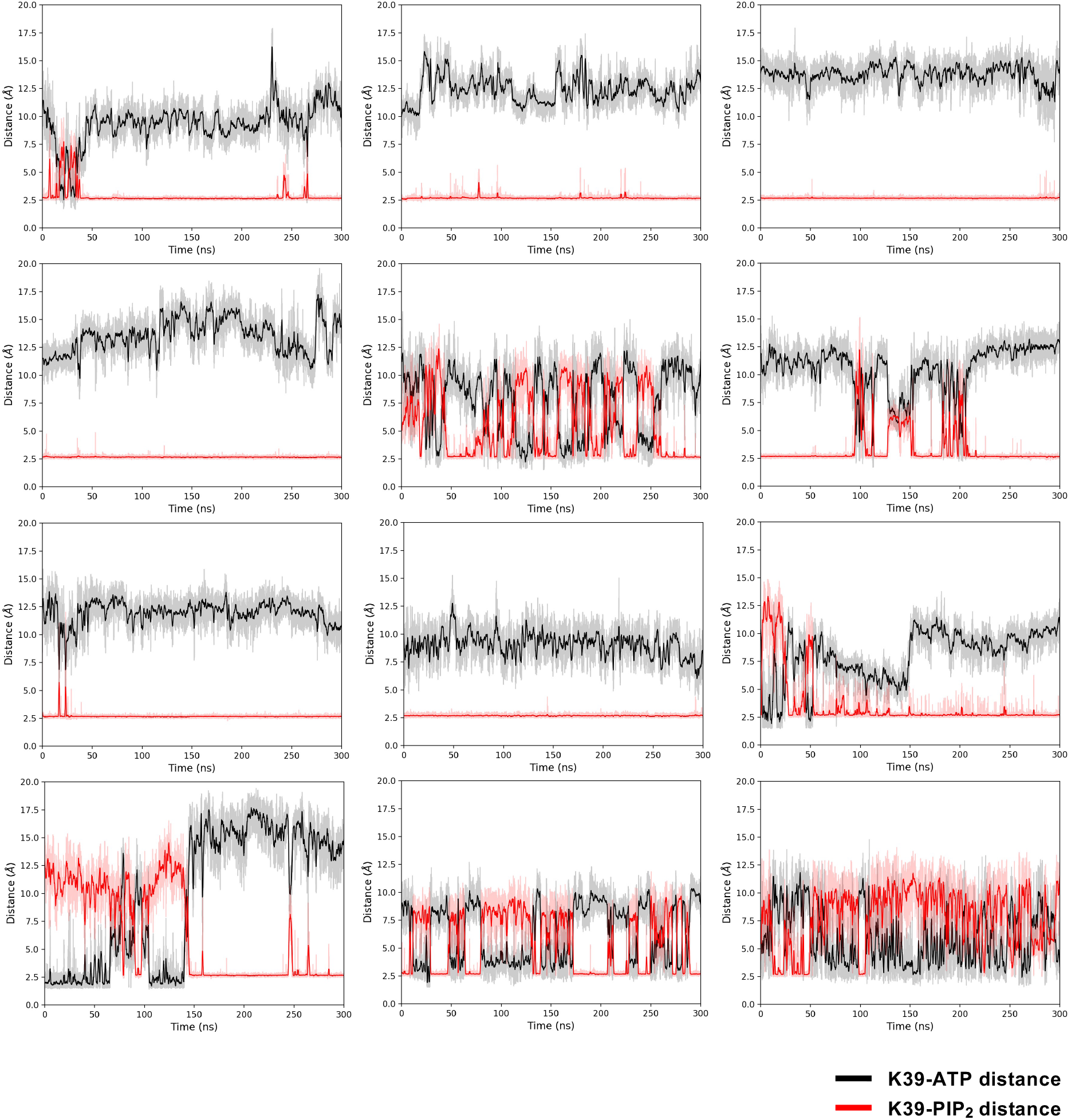
The distance between K39, ATP and PIP_2_. Distance calculation of the ATP (black) and PIP_2_ (red) and the K39 when both ligands are present, measured over the final 300 ns simulation trajectory. The darker lines show running averages of every 1 ns of the simulation.

**Supplementary figure 4.**
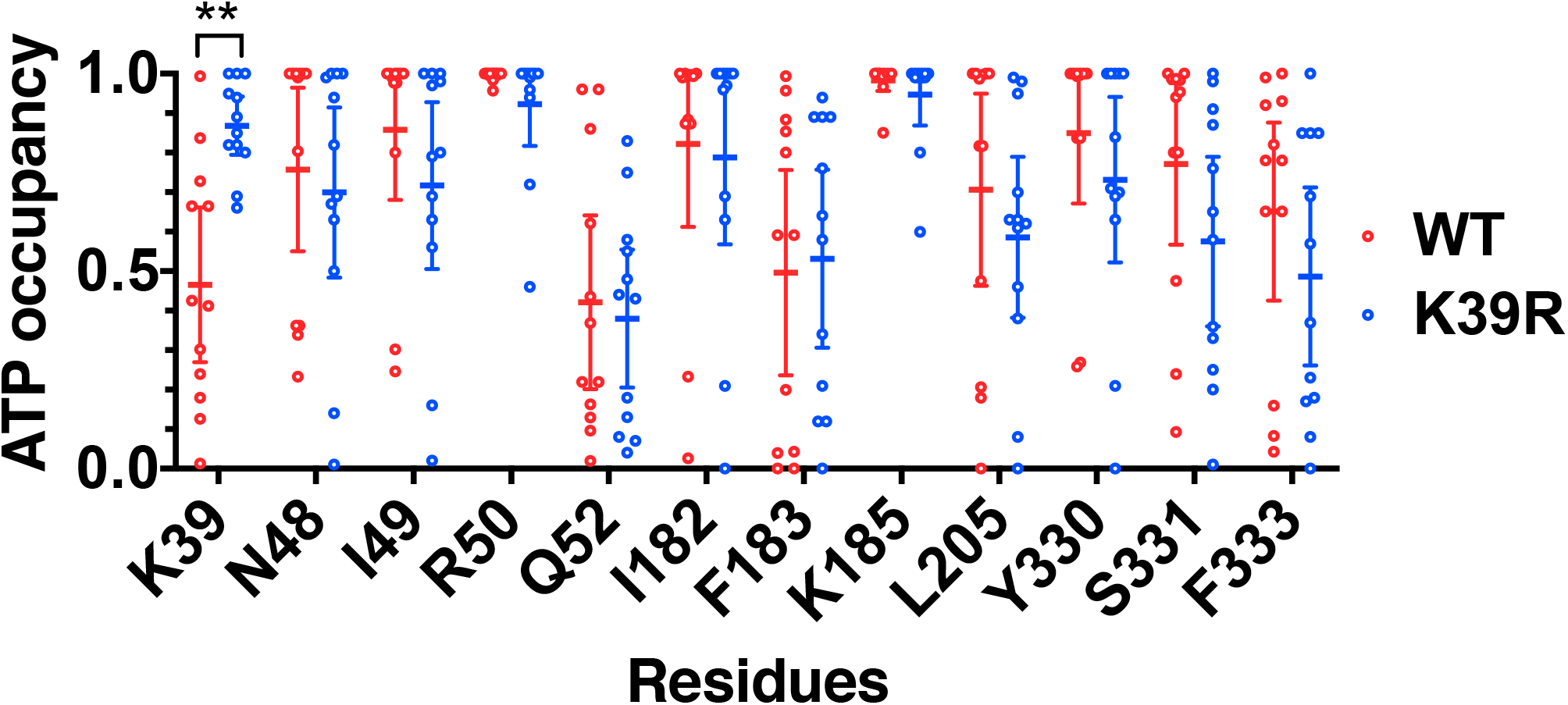
An effect of ATP occupancy caused by K39R mutation. ATP contact analysis showing the fraction of time that residues which contact the ATP molecule >40% in all 12 repeats of 300 ns simulations are within 4 Å proximity to the channel (ATP occupancy) for WT (red) and the K39R mutant (blue). The error bar indicates the 95% confidence interval around the mean. ** p < 0.01 (Student’s t-test).

**Supplementary figure 5.**
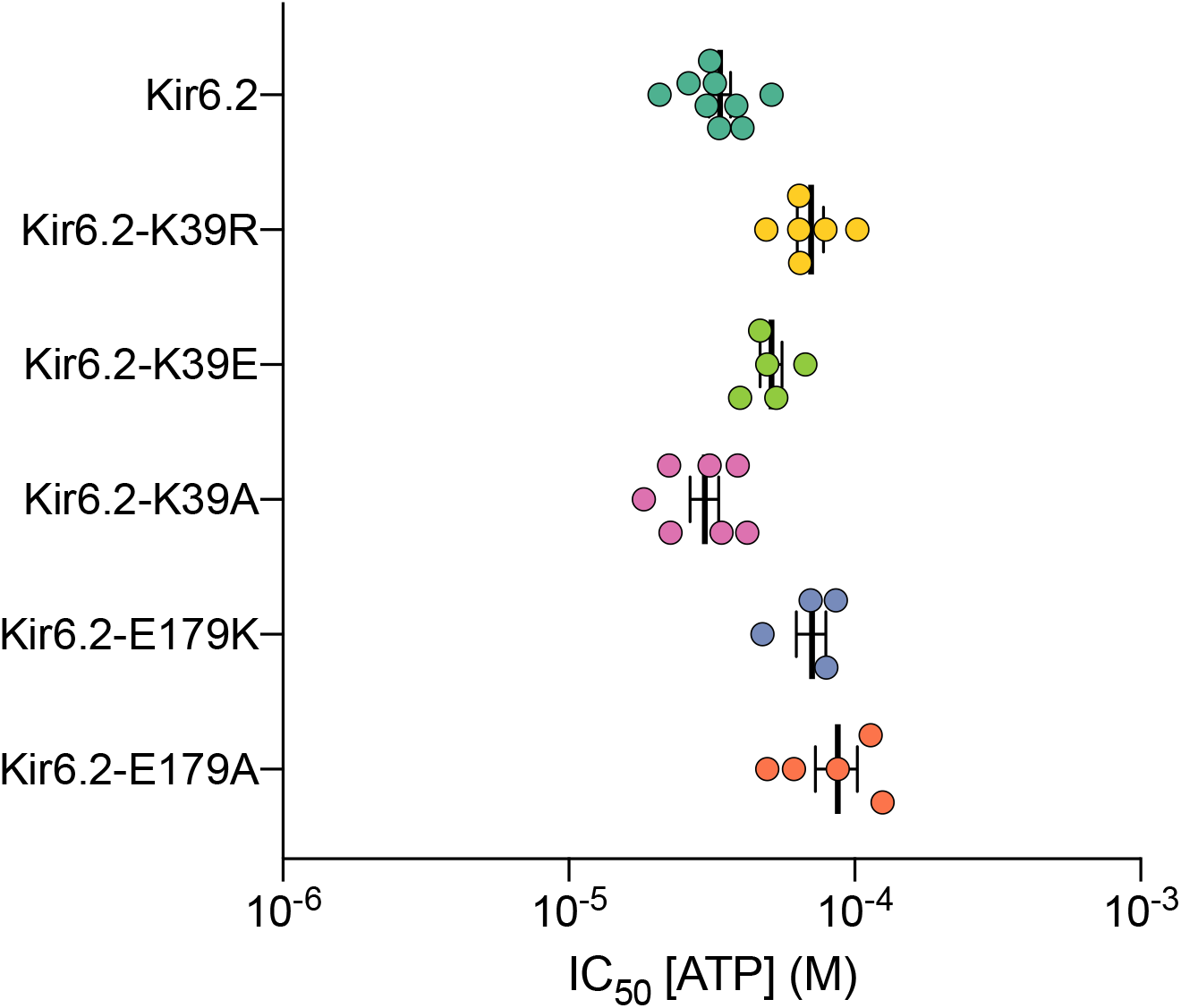
Effect of ATP on K_ATP_ current. IC_50_s from ATP current inhibition derived from Hill fits to individual experiments are plotted as coloured data points. The means and 95% confidence intervals for each construct are overlaid as black error bars. Each construct is labelled with GFP at the C-terminus, but not with ANAP at residue W311.

**Supplementary figure 6.**
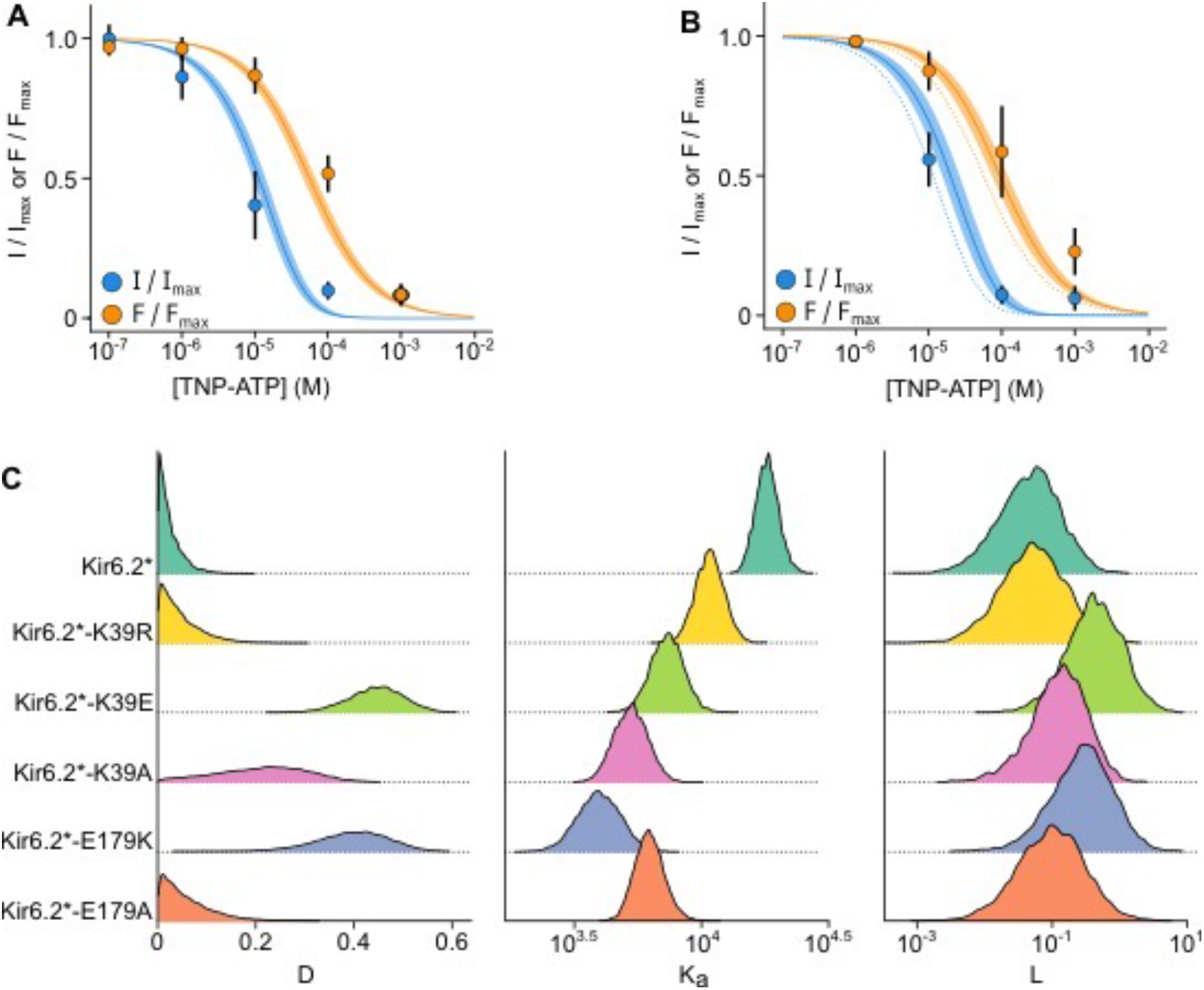
MWC-type modelling. A, B) Concentration-response relationships for TNP-ATP inhibition of currents (I/I_max_) and for quenching of ANAP fluorescence (F/F_max_) for Kir6.2* (A) or Kir6.2*-K39R (B). Data points are the mean and error bars are the 95% confidence intervals. Data were fit to an MWC-type model. Solid curves represent the median fits and shaded areas indicate the 95% quantile intervals. B) The median fits to Kir6.2* in panel A are shown here as dotted lines. C) Posterior probability distributions for the MWC-type model parameters for each construct.

**Supplementary figure 7.**
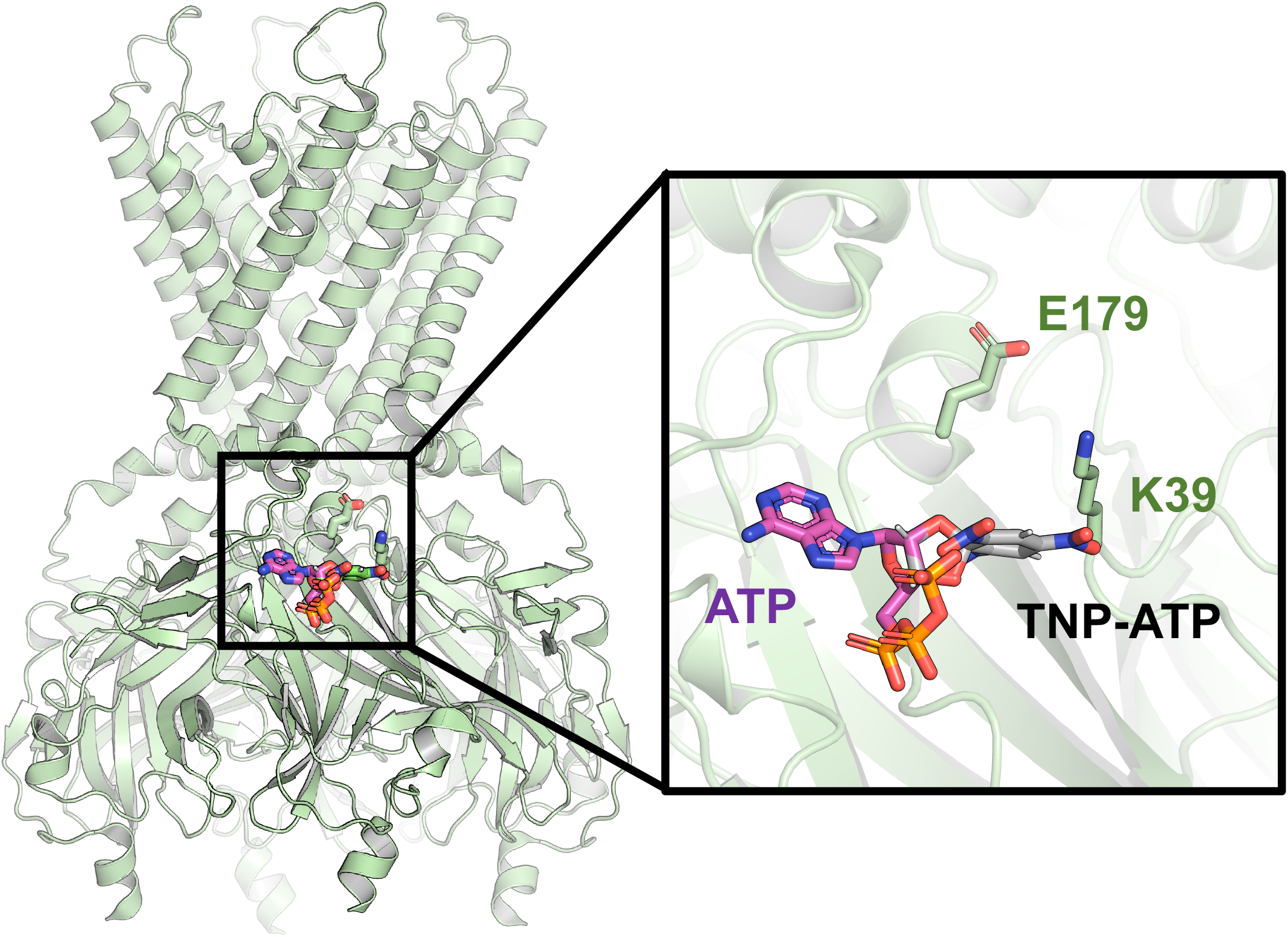
Alignment of TNP-ATP and ATP on the Kir6.2 structure. Alignment of TNP-ATP to ATP on the Kir6.2 structure. TNP-ATP is represented in grey, ATP in purple. K39 and E179 side chains are represented as stick in green.

**Supplementary table 1.** IC_50_s from ATP current inhibition of Kir6.2 with SUR1. Each construct is labelled with GFP at the C-terminus, but not with ANAP at residue W311. The number of repeats per construct is shown in parenthesis. The error shows standard error of the mean. *h* = 1.3-1.7 for fits to the concentration-response relationships for ATP inhibition. p-values are calculated using Dunnet’s post-hoc test in comparison to control (Kir6.2) after a one-way ANOVA.

| Construct | IC<sub>50</sub> (μM)<br>[ATP] | Dunnett's adjusted p-value |
| --- | --- | --- |
| Kir6.2 | 34±3 (9) | Control |
| Kir6.2-K39A | 31±4 (7) | 0.95 |
| Kir6.2-K39E | 61±7 (5) | <0.01 |
| Kir6.2-K39R | 74±7 (6) | <0.01 |
| Kir6.2-E179A | 90±12 (5) | <0.01 |
| Kir6.2-E179K | 82±8 (4) | <0.01 |

**Supplementary table 2.** IC_50_s from ATP current inhibition of Kir6.2* with SUR1. Each construct is labelled with GFP at the C-terminus, and ANAP labelled at residue W311. The number of repeats per construct is shown in parenthesis. The error shows standard error of the mean. *h* = 1.3-1.7 for fits to the concentration-response relationships for ATP inhibition. p-values are calculated using Dunnet’s post-hoc test in comparison to control (Kir6.2*) after a one-way ANOVA.

| Construct | IC<sub>50</sub> (μM)<br>[ATP] | Dunnett's adjusted p-value |
| --- | --- | --- |
| Kir6.2* | 112±9 (8) | Control |
| Kir6.2*-K39A | 169±21 (8) | 0.15 |
| Kir6.2*-K39E | 173±22 (6) | 0.038 |
| Kir6.2*-K39R | 445±33 (8) | <0.01 |
| Kir6.2*-E179A | 376±80 (5) | <0.01 |
| Kir6.2*-E179K | 351±63 (8) | <0.01 |

**Supplementary table 3.** IC_50_s from ATP current inhibition of Kir6.2 without SUR1. Each construct is labelled with GFP at the C-terminus, but not with ANAP at residue W311. The number of repeats per construct is shown in parenthesis. *nd*, not determined*, no macroscopic current was measured when mutant was expressed in the absence of SUR1. *h* = 0.9-1.3 for fits to the concentration-response relationships for ATP inhibition. The error shows standard error of the mean.

| Construct | IC<sub>50</sub> (μM)<br>[ATP] |
| --- | --- |
| Kir6.2 | 208±36 (3) |
| Kir6.2-K39A | *nd* * |
| Kir6.2-K39E | 1280±na (1) |
| Kir6.2-K39R | 384±19 (3) |
| Kir6.2-E179A | 1293±304 (4) |
| Kir6.2-E179K | 378±178 (4) |

**Supplementary table 4.** IC_50_s from TNP-ATP current inhibition of Kir6.2 with SUR1. Each construct is labelled with GFP at the C-terminus, but not with ANAP at residue W311. Brackets show number of repeats per construct. The error shows standard error of the mean. *h* = 1.3-1.7 for Hill fits to the concentration-response relationships of individual experiments for TNP-ATP inhibition p-values are calculated using Dunnet’s post-hoc test in comparison to control (Kir6.2) after a one-way ANOVA.

| **Construct** | **IC<sub>50</sub> (μM)**<br>**[TNP-ATP]** | **Dunnett's adjusted**<br>**p-value** |
| --- | --- | --- |
| **Kir6.2** | 1.39±0.40 (12) | Control |
| **Kir6.2-K39A** | 13.2±0.28 (8) | <0.01 |
| **Kir6.2-K39E** | 12.6±0.44 (5) | <0.01 |
| **Kir6.2-K39R** | 2.62±0.44 (9) | 0.054 |
| **Kir6.2-E179A** | 13.6±3.24 (7) | <0.01 |
| **Kir6.2-E179K** | 56.3±17.8 (7) | <0.01 |

**Supplementary table 5.** IC_50_s from TNP-ATP current inhibition of Kir6.2* with SUR1. Each construct is labelled with GFP at the C-terminus, but with ANAP labelled at residue W311. Brackets show number of repeats per construct. The error shows standard error of the mean. *h*= 0.9-1.3 for Hill fits to the concentration-response relationships of individual experiments of TNP-ATP inhibition p-values are calculated using Dunnet’s post-hoc test in comparison to control (Kir6.2*) after a one-way ANOVA.

|  | **IC<sub>50</sub> (μM)**<br>**[TNP-ATP]** | **Dunnett's adjusted p-**<br>**value** |
| --- | --- | --- |
| **Kir6.2\*** | 7.06±1.79 (11) | Control |
| **Kir6.2\*-K39A** | 46.8±9.48 (12) | <0.01 |
| **Kir6.2\*-K39E** | 110±45.2 (8) | <0.01 |
| **Kir6.2\*-K39R** | 9.81±0.90 (6) | 0.44 |
| **Kir6.2\*-E179A** | 27.7±5.23 (6) | <0.01 |
| **Kir6.2\*-E179K** | 169±61.7 (7) | <0.01 |

**Supplementary table 6.** EC_50_s from TNP-ATP FRET of Kir6.2* with SUR1. Each construct is labelled with GFP at the C-terminus, but with ANAP labelled at residue W311. Brackets show number of repeats per construct. The error shows standard error of the mean. *h* = 0.9-1.3 for Hill fits to the concentration-response relationships of individual experiments of TNP-ATP inhibition. p-values are calculated using Dunnet’s post-hoc test in comparison to control (Kir6.2*) after a one-way ANOVA.

|  | **EC<sub>50</sub> (μM)**<br>**[TNP-ATP]** | **Dunnett's adjusted p-value** |
| --- | --- | --- |
| **Kir6.2\*** | 94.4±14.0 (11) | Control |
| **Kir6.2\*-K39A** | 244±61.4 (11) | <0.01 |
| **Kir6.2\*-K39E** | 165±60.1 (8) | 0.02 |
| **Kir6.2\*-K39R** | 209±72.7 (6) | <0.01 |
| **Kir6.2\*-E179A** | 170±23.6 (6) | <0.01 |
| **Kir6.2\*-E179K** | 171±60.5 (7) | <0.01 |

